# Potential Neurocognitive Biomarkers for Post Traumatic Stress Disorder (PTSD) Severity in Recent Trauma Survivors

**DOI:** 10.1101/721068

**Authors:** Ziv Ben-Zion, Yoav Zeevi, Nimrod Jackob Keynan, Roee Admon, Tal Kozlovski, Haggai Sharon, Pinchas Halpern, Israel Liberzon, Arieh Y. Shalev, Yoav Benjamini, Talma Hendler

## Abstract

Contemporary symptom-based diagnosis of Post-traumatic Stress Disorder (PTSD) largely overlooks related neurobehavioral findings and rely entirely on subjective interpersonal reporting. Previous studies associating objective biomarkers with PTSD have mostly used the disorder’s symptom-based diagnosis as main outcome measure, overlooking the actual clustering and richness of phenotypical features associated with PTSD. Here, we aimed to computationally derive potential neurocognitive biomarkers that could efficiently differentiate PTSD subtypes, based on an observational cohort study of recent trauma survivors. A three-staged semi-unsupervised method (“3C”) was used to *categorize* trauma survivors based on current PTSD diagnostics, derive *clusters* of PTSD based on features related to symptom load, and to *classify* participants’ cluster membership using objective features. A total of 256 features were extracted from psychometrics, cognitive, structural and functional neuroimaging data, obtained from 101 adult civilians (age=34.80±11.95, 51 females) evaluated within a month of trauma exposure. Multi-domain features that best differentiated cluster membership were indicated by using importance analysis, classification trees, and ANOVA. Results revealed that entorhinal and rostral anterior cingulate cortices volumes (structural domain), in-task amygdala’s functional connectivity with the insula and thalamus (functional domain), executive function and cognitive flexibility (cognitive domain) best differentiated between two clusters related to PTSD severity. Cross-validation established the results’ robustness and consistency within this sample. Multi-domain biomarkers revealed by the 3C analytics offer objective classifiers of post-traumatic morbidity shortly following trauma. They also map onto previously documented neurobehavioral PTSD features, supporting the future use of standardized and objective measurements to more precisely identify psychopathology subgroups shortly after trauma.

## Introduction

Post-traumatic stress disorder (PTSD) symptoms are commonly observed shortly after exposure to trauma, with the extent of their initial intensity associated with a high risk for poor-recovery^1–3^. Categorization is currently based on symptoms reported via structured interviews, such as the Clinician-Administered PTSD Scale (CAPS). While such scales show good reliability, construct and predictive validity, they also have notable limitations. The CAPS does not capture various symptoms often co-expressed with PTSD, such as depression and general anxiety^4^. It is also only weakly linked with cognitive measures and other objective measures of putative biological mechanisms^5–7^. By solely accounting for reported- or observed symptoms, the CAPS entirely rely on subjective interpersonal reporting. These limitations may be responsible for its long term diagnoses instability ^8^ as well as its suboptimal use as a guide for individualized clinical management^9^.

*Cognitive functioning* is one of many dimensions often overlooked in the clinical context of post-traumatic psychopathology. To date, reported associations between cognitive deficits and PTSD included working memory, information processing speed, verbal learning, short-term and declarative memory, attention and executive functioning, response inhibition and attentional switching (see studies reported in recent meta-analyses^10,11^). Adaptive cognitive functioning has been linked with resilience and a reduced likelihood of post-traumatic stress symptoms development and maintenance^11,12^. The investigation of *brain structure and function* are also crucial for understanding the underlying clinical manifestations of PTSD. Indeed, accumulating evidence from neuroimaging points to structural and functional brain abnormalities among PTSD patients^5,13–16^, including lower hippocampal volume^17–19^ and altered activity and connectivity of the amygdala, insula, ACC, mPFC and dlPFC; structures known to be involved in threat detection, executive function, emotion regulation, and contextual processing^5,13,20–24^. Some structural and functional neuroimaging studies also point to early predisposing factors in the development of PTSD^13,21,25–27^. Despite promising findings from cognitive and neuroimaging studies, these objective measures have yet to be integrated in the routine assessment and management of post-traumatic psychopathology. One obstacle in the clinical translation of these findings is an unclear understanding of how these indicators cluster together into PTSD subtype to provide a better guide for potential interventions. Moreover, most studies associating objective biomarkers with PTSD have used the disorder’s symptom-based formal diagnostic criteria as main outcome measure, thereby overlooking the actual clustering and richness of phenotypical features associated with post-traumatic psychopathology within a dataset, and imposing an extraneous construct (PTSD diagnosis) as an outcome of interest.

The above-mentioned limitations could benefit from advanced data-driven computational and statistical methods that can evaluate wider arrays of potential biomarkers and disorder indicators, quantifying their relationship with various clinical manifestations. Machine learning methods are particularly well-suited to confront such computational challenges, as they are able to account for the complex interrelation of many relevant factors^28^. Indeed, the last decade has shown an exponential increase in the use of machine learning for the study of posttraumatic stress, including both supervised and unsupervised approaches. Supervised analytic approaches are limited by the accuracy of the prior knowledge they rely on^29^. While unsupervised approaches make few to no assumptions, this limits the subpopulations revealed, as they are not tied to any specific questions of interest^29^.

A possible solution is the use of hybrid analytic methods that combine both supervised and unsupervised approaches, which may lead to more precise diagnosis by identifying novel combinations of potential biomarkers for specific disorders^29^. The present study propose to evaluate objective biomarkers’ ability to identify data-driven clusters, which hopefully represent post-traumatic psychopathology phenotypes. To do so, we applied a recently developed three-staged hybrid analytic methodology, termed 3C (Categorize, Cluster, and Classify); a semi-unsupervised method that combines theory- and data-driven approaches^30^. Analysis included both clinical wide-used knowledge assumptions and state-of-the-art data-driven methods apply on objective measures obtained from neuroimaging and cognitive testing of recent trauma survivors. We assumed that such a hybrid analytic approach would unveil a unique set of potential mechanism-related neurocognitive biomarkers for PTSD, closely tied to pre-existing diagnostic methods. The term “subtypes” accounts for different demographics, clinical sub-scales or symptom severity.

## Materials and Methods

The 3C methodology was used on a multi-domain data set composed of clinical interviews and questionnaires, computerized cognitive testing and neuroimaging of brain structure and function. Data was obtained from recent trauma survivors discharged from a general hospital’s emergency department (ED) after experiencing a traumatic event. This data is a part of a larger on-going project that examines PTSD development in trauma survivors (for full study protocol please see Ben-Zion et al. (2019)^31^). Here we present multi-domain dataset obtained from participants within one-month of the traumatic incident.

### Participants

101 recent trauma survivors (age=34.80±11.95, range 18-65, 51 females) admitted to ED following a traumatic experience. The most common trauma types were motor-vehicle accidents (n=79, 78%), and other traumatic events included assaults, terror attacks, hostilities, drowning, mass casualty incidents, robbery and electrocution. Out of 101 participants, n=58 individuals met a full PTSD symptom diagnosis during the face-to-face clinical interview (“PTSD” group), and n=43 did not meet this criterion even though they still suffered from a variety of PTSD symptoms (“No PTSD” group).

### Clinical Instruments

PTSD symptom severity was quantified using the Clinician-Administered PTSD Scale for DSM-IV (CAPS-4)^32^. Four additional self-report clinical questionnaires were administered: PTSD Checklist for DSM-IV (PCL-4)^33^ evaluating post-traumatic symptoms; Beck Depression Inventory (BDI)^34^ assessing current depressive symptoms; Anxiety Inventory (BAI)^35^ measuring concurrent anxiety symptoms; and Participants’ Clinical Global Impression Scale (CGI-P)^36^ evaluating patients’ subjective impression, on a scale between 1 (‘normal feeling’) to 7 (‘the worst feeling there is’). For more details on these clinical instruments, please see^31^.

### Cognitive Functioning

WebNeuro^37^, an internet-based comprehensive cognitive assessment battery previously validated against traditional cognitive tests, was used to assess cognitive functioning. To standardize testing conditions, all tests were conducted at our laboratory in Hebrew. Performance on the different tasks was calculated using an automated software program that derived standardized Z-scores for each participant on each of the following eleven cognitive domains: motor coordination, processing speed, sustained attention, controlled attention, cognitive flexibility, response inhibition, working memory, recall memory, executive function, emotion identification, and emotional bias. For detailed description of the cognitive tasks, please see^12,31^.

### Imaging Data Acquisition

Structural and functional scans were performed in a 3.0 Tesla Siemens MRI system (MAGNETOM Prisma, Germany), using a twenty-channel head coil, located in our lab at Tel-Aviv Sourasky Medical Center. To allow high-resolution whole-brain structural images, a T1-weighted magnetization prepared rapid gradient echo (MPRAGE) (TR/TE=2400/2.29ms, flip angle = 8°, voxel size 0.7 × 0.7 × 0.7 mm, FOV = 224 × 224 mm) was acquired. Functional whole-brain scan was performed in an interleaved order, using a T2*-weighted gradient echoplanar imaging pulse sequence (TR/TE = 2000/28 ms, flip angle = 90°, voxel size 2.2 × 2.2 × 2.2 mm, FOV =220 × 220 mm, slice thickness = 3mm, 36 slices per volume).

### Structural Imaging Data Analysis

Cortical reconstruction and volumetric segmentation were performed with the FreeSurfer image analysis suite^38^ (version 1.379.2.73), which is documented and freely available for download online (http://surfer.nmr.mgh.harvard.edu/). Briefly, this processing included motion correction and the averaging^39^ of multiple volumetric T1 weighted images, removal of non-brain tissue using a hybrid watershed/surface deformation procedure^40^, automated Talairach transformation, segmentation of the subcortical white matter and deep gray matter volumetric structures (including hippocampus, amygdala, caudate, putamen, ventricles)^41,42^, intensity normalization, tessellation of the gray matter-white matter boundary, automated topology correction, and surface deformation following intensity gradients to optimally place the gray/white and gray/cerebrospinal fluid borders at the location where the greatest shift in intensity defines the transition to the other tissue class. The automatic subcortical segmentation of brain volume is based upon the existence of an atlas (“aseg”) containing probabilistic information on the location of structures^43^. The maps are created using spatial intensity gradients across tissue classes and are therefore not simply reliant on absolute signal intensity. The maps produced are not restricted to the voxel resolution of the original data thus are capable of detecting submillimeter differences between groups.

### Functional Imaging Data Analysis

Preprocessing and statistical analysis of the functional images were performed in a voxel-based approach using Statistical Parametric Mapping (SPM)^44^ version 12. In short, this process included: Slice time correction, using one slice before the last as the reference slice. Head motions correction by six-parameter rigid body spatial transformations, using three translations and three rotation parameters, with the first image serving as a volume reference. A 4^th^ degree interpolation was applied to detect and correct head motions. Functional maps were automatically co-registered to corresponding structural maps using an objective function of normalized mutual information (NMI). The complete dataset was transformed into MNI space and spatially smoothed with an isotropic 6mm full-width at half-maximum (FWHM) Gaussian kernel.

During this scan, participants performed the **Emotional Faces Matching Task**^45^, which was used to evaluate their emotional reactivity. In this task, subjects were instructed to select the face/shape (located at the bottom right or bottom left of the screen) that matches the target face/shape (located at the top of the screen), as accurately and as quickly as possible. The tasks included four blocks of shapes (that were used as a baseline) and four blocks of emotional faces (angry, fearful, surprised and neutral faces). The order of the blocks of emotional faces was counterbalanced between subjects using four different versions for this task. Both whole-brain activations and functional connectivity of the amygdala (seed region) were calculated for the following contrasts: angry faces (vs. shapes), fearful faces (vs. shapes), surprised faces (vs. shapes), and neutral faces (vs. shapes). For a full list of the functional brain measures derived from this analysis, please refer to ‘brain function variables’ in Supplementary Table 1.

### Procedure

A member of the research team identified potentially trauma-exposed patients using the ED medical records. Within 10–14 days after potential trauma exposure, and after being discharged from the hospital, these individuals were contacted for an initial telephone screening, which was conducted by MA-level clinicians that were trained in the specific assessment tools. After obtaining verbal consent, the PCL-5 was administered to assess the risk of PTSD development. Those who met PTSD symptom criteria (except the “one-month duration” criteria) and did not meet any of the exclusion criteria (see under Participants section), received verbal information about the study. Participants were subsequently invited to participate in two meetings, both within 30 days from ER admission: (1) In-person clinical assessment including administration of the CAPS, self-report questionnaires (BDI, BAI, PCL, CGI) and WebNeuro cognitive battery; (2) Structural and functional MRI scan. Each meeting lasted approximately 3 hours (total of 6 hours for both), for which participants received financial remuneration, in accordance with the ethics committee regulations and approval.

### Statistical Approach

The 3C method^30^ assumes that existing medical knowledge of a given disorder is critical but not sufficient for an accurate diagnosis. It suggests to build upon the current diagnostics and expands upon it with unsupervised data-driven methods. Thus, using well-used clinical measures to divide patients into homogeneous clusters based on common characteristics. Clustering allows exploration of objective measures related to the specific disorder-subtype. A recent study using the 3C^46^ discovered new sub-phenotypic groups and their specific biomarkers in a large Alzheimer’s disease dataset (“ADNI”^47^). The 3C finds homogeneous groups via unsupervised clustering using well-used clinical measures, relevant to the disease diagnosis, it then further characterizes those groups by exploring potential biomarkers. The 3C procedure is composed of three stages; Categorize, Cluster and Classify. During <u>Categorization</u>, multi-domain variables are sorted into three categories; 1) *Assigned diagnosis*; as currently applied in the field. 2) *Clinical measuremen*t; variables that describe the patient’s condition and the expression of the disease (symptoms and signs). 3) *Potential biomarkers*; variables that could improve existing diagnostic procedures, but not currently in use in the diagnostic procedure. <u>Clustering</u> includes two steps: 1) A supervised selection of the most relevant *clinical variables* through some statistical testing, to be used as predictors to the *assigned diagnosis*. This feature selection was done based on a permissive threshold of Benjamini-Hochberg^48^ FDR-adjusted of p=0.2 (From now on, the use of “FDR” in this manuscript will specifically refer to the BH procedure). This was done to eliminate clinical variables that are not relevant to the disease as defined by *the assigned diagnosis* category. 2) Unsupervised clustering utilizing the selected *clinical measurements* was performed, using k-medoids with Manhattan distance metrics. This allowed us to discover data-driven homogeneous clusters that are related to existing diagnostics (i.e., dividing participants into subtypes based on common–used variables). Nevertheless, these clusters are not limited to formal symptom-based PTSD diagnosis (as indicated by CAPS), but rather capture the actual richness of phenotypical features associated with post-traumatic psychopathology within a dataset. Prior to clustering we determined the optimal number of clusters based on two metrics: Gap statistics^49^ and Silhouette^50^. Lastly, <u>Classification</u> includes characterization of the clusters based on the objective *potential biomarkers*, via three distinct approaches: importance analysis (mean decrease GINI^51^); classification trees; and a marginal one-way analysis of variance (ANOVA) between clusters for each potential bio-marker.

### Algorithms, Codes, and Software

Algorithms and codes were performed using R software version 3.4.4. Data imputation was performed using the 5-nn method in order to deal with missing data (less than 1% of the full dataset). Variables were monotonic transformed to gain symmetry when needed, using a semi-automated Shiny App^52^. Importance, measured as the marginal loss of classification accuracy for each variable by randomly permuting it on the out-of-bag validation set, was calculated using the {randomForest} R package^51^. R package {cluster}^53^ was used for clustering, and R package {rpart} was used for classification and regression trees (CART).

## Results

During <u>Categorization</u>, features were divided into three distinct categories required by the 3C method: *Assigned Diagnosis* was based on the CAPS-4 total scores, *Clinical Measurements* included the total scores of the four self-report questionnaires (PCL, BDI, BAI, and CGI), and *Potential Biomarkers* included 11 standardized total scores obtained from computerized cognitive testing, 192 features from structural imaging (including volumes and thickness of subcortical and cortical areas), 48 features extracted from fMRI during the emotional faces matching task (including whole-brain activations and functional connectivity of left and right amygdala during the task). For a full list of neurocognitive features, see Supplementary Table 1.

For <u>Clustering</u>, we used the *Clinical Measurements* that were found highly correlated with PTSD symptom severity as indicated by CAPS-4 total scores: PCL, BDI, BAI, and CGI (based on the 0.2 FDR threshold). Participants were divided into an optimal number of two clusters, based on both Gap statistics^49^ and Silhouette^54^ methods, which presented the best separation on all four clinical measurements (PCL, BDI, BAI, and CGI). Participants within the first cluster showed an average higher severity on all four clinical measurements, compared to those within the second cluster (see Fig. 1). To test the association between these clusters (representing high and low “disease load”) and PTSD diagnosis (PTSD or not according to CAPS-4), a two-sample test for equality of PTSD proportions between the two clusters was conducted. Results revealed a significant link between the proposed clusters in PTSD dichotomous diagnosis (*F*_*1,99*_*=35.47*, *p<0.001*, *⍰*^*2,1*^*=20.911*, *p<0.001*, *CI of the difference of proportions = (0.28,0.63)*) (see Table 1). Furthermore, a one-way ANOVA was performed on the CAPS-4 total scores (continuous PTSD symptom severity measure) with clusters as the group variable. Results indicated that individuals in cluster 2 had significantly higher CAPS-4 total scores *(M=61.91±18.16)* compared to those in cluster 1 *(M=37.45±23.12)(p<0.001)* (see Supplementary Figure 1). It is important to note that these clusters do not represent a simple division into high and low CAPS severity scores, but rather a data-driven division into severity subtypes, based on self-reported symptoms of depression (BDI), anxiety (BAI), post-trauma (PCL) and the patients’ subjective global impression (CGI). As mentioned, these subtypes (clusters) were found to correlate with PTSD clinical diagnosis and severity, but were not identical to it, and rather represent clusters of “disease load”. Accordingly, cluster 1 will now be referred to as the “Low-Symptomatic” Cluster (LoClus, low PTSD severity), and cluster 2 as the “High-Symptomatic” Cluster (HiClus, high PTSD severity).

**Table 1.**
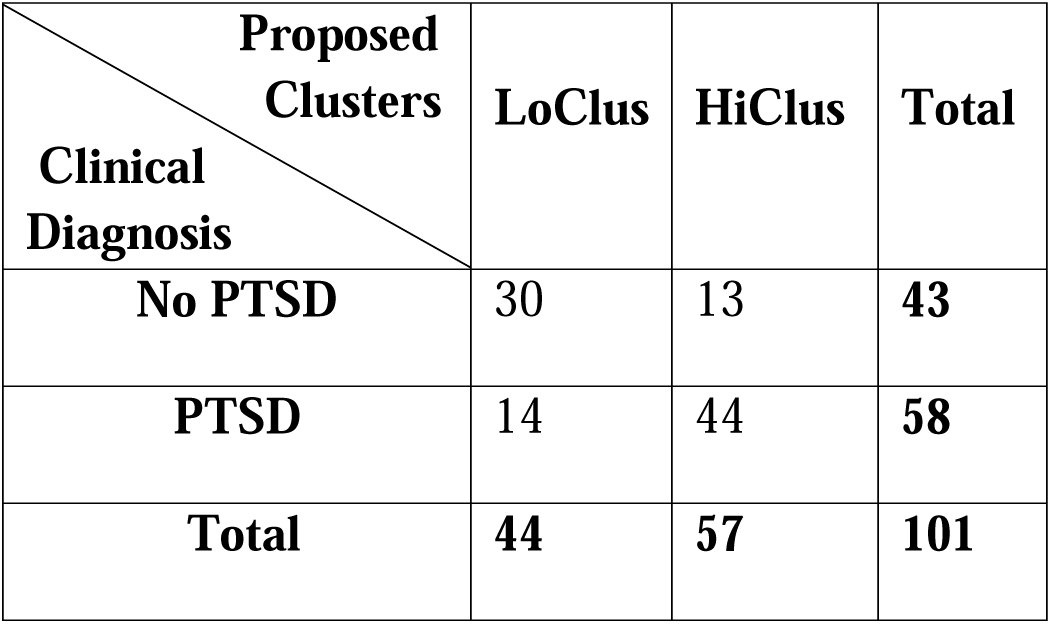
Confusion matrix of PTSD diagnosis clusters assignments. Table rows represent individuals’ current clinical DSM-based PTSD diagnosis (PTSD / No PTSD), while the columns represent the two proposed clusters (LoClus=Low-Symptomatic Cluster / HiClus=High-Symptomatic Cluster).

| **Clinical**<br>**Diagnosis \ Proposed**<br>**Clusters** | **LoClus** | **HiClus** | **Total** |
| --- | --- | --- | --- |
| **No PTSD** | 30 | 13 | **43** |
| **PTSD** | 14 | 44 | **58** |
| **Total** | **44** | **57** | **101** |

**Figure 1.**
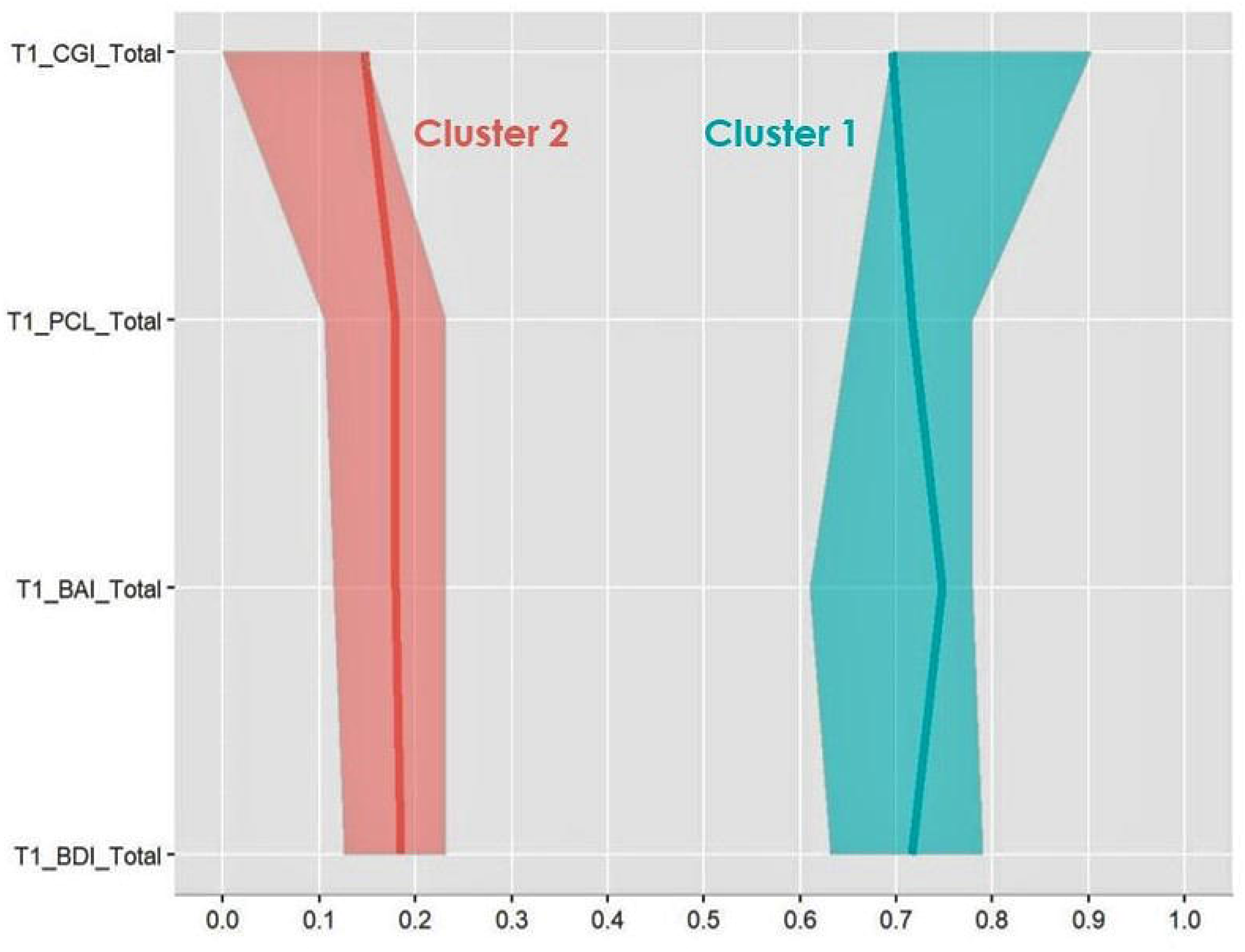
Parallel Coordinates Plot of Clinical Measurements. The Y-axis depicts the four variables on which the clusters were built, all of them with an FDR-adjusted p-value smaller than 0.2, while the x-axis depicts their percentiles. The figure presents, for each cluster, its median value (bold line) and their 0.025 and 0.975 percentiles (i.e., 95% CI, the scattered “cloud” around it) of 400 bootstrap samplings’ medians. Participants belonging to the first cluster (turquoise) were, on average, higher on all four variables, compared to those belonging to the second cluster (red). “T1_CGI_Total” = Total Score of Clinical Global Impression Scale Questionnaire; “T1_PCL_Total” = Total Score of PTSD Checklist Questionnaire; “T1_BAI_Total” = Total Score of Beck Anxiety Inventory Questionnaire; “T1_BDI_Total”= Total Score of Beck Depression Inventory Questionnaire.

During the <u>Classification</u> stage, clusters were based on objective variables that could serve as potential biomarkers, using a mean decrease GINI index (i.e. importance index) for each biomarker. The most significant potential neurocognitive biomarkers associated with the resulted clusters included left EC volume *(Importance=0.884)*, cognitive flexibility *(Importance=0.487)*, rostral anterior cingulate cortex (rACC) volume *(Importance=0.429)*, and average amygdala functional connectivity with the thalamus while watching angry faces vs. shapes (Importance=0.419). The top ten potential neurocognitive biomarkers for the clustering are presented in Figure 2 and Supplementary Table 2.

**Figure 2.**
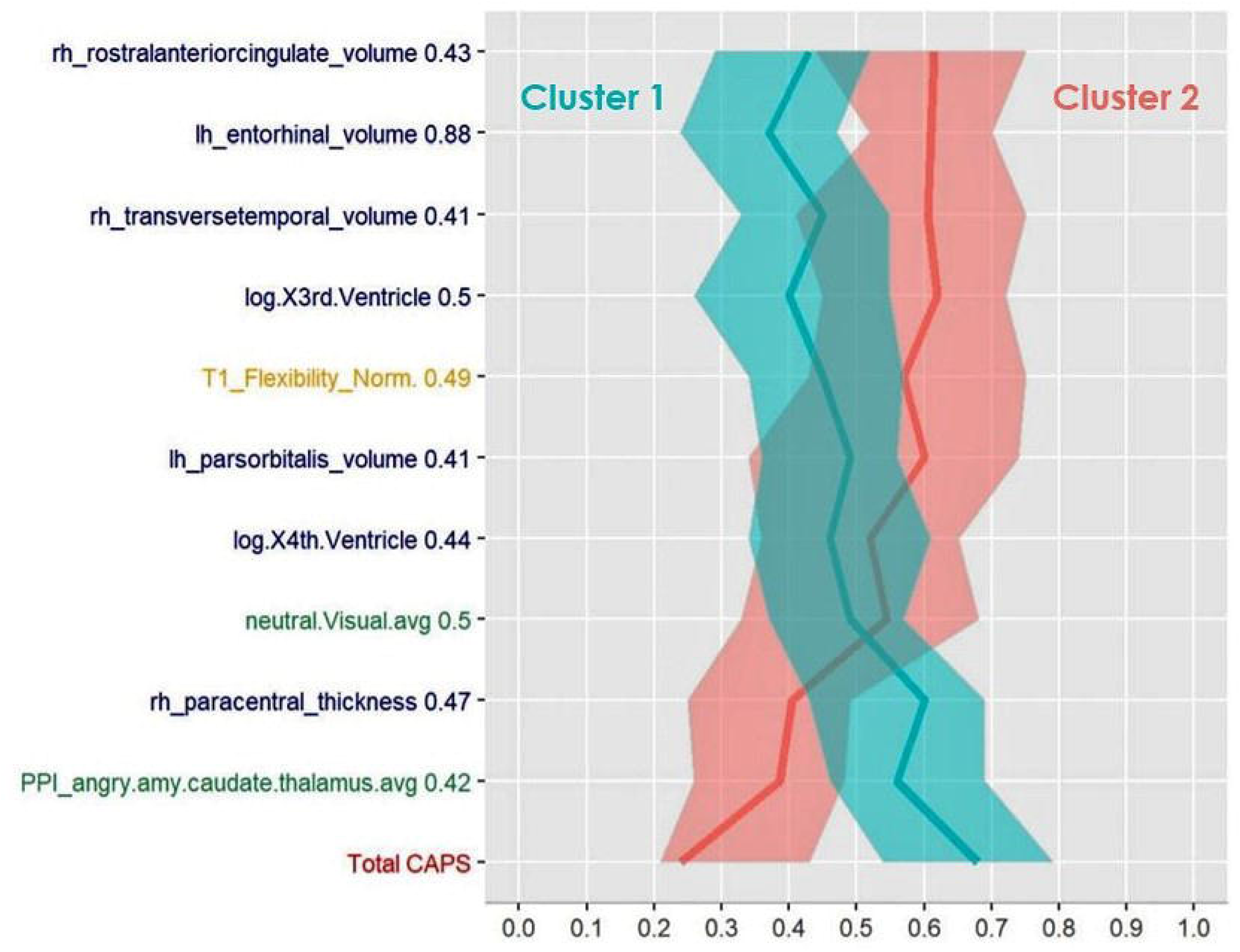
Parallel Coordinates Plot of Potential Neurocognitive Biomarkers. The Y-axis depicts the top ten most important potential neurocognitive biomarkers in classifying to the two clusters, together with their mean decrease GINI measure (i.e. importance index). Biomarkers from cognitive domains are colored in yellow, biomarkers from structural brain measurements are colored in dark blue, and biomarkers from functional brain measurements are colored in green. Average CAPS-4 total score is presented for both “Low-Symptomatic” Cluster (cluster 1, turquoise) and “High-Symptomatic” Cluster (cluster 1, red). The medians of 400 Bootstrap samplings were drawn, and their median and 0.025 and 0.975 quantiles are plotted per cluster. “rh_rostralanteriorcingulate_volume” = right rostral anterior cingulate cortex volume; “lh_entorhinal_volume” = left entorhinal cortex volume; “rh_transversetemporal_volume” = right transverse temporal gyrus volume; “log.X3rd.Ventricle” = third ventricle volume (after log transformation to create symmetrical distribution); “T1_Flexibility_Norm” = cognitive flexibility score; “lh_parsorbitalis_volume” = left pars orbitalis volume; “log.X4th.Ventricle” = fourth ventricle volume (after log transformation to create symmetrical distribution); “neutral.Visual.avg” = average visual cortex activity in response to neutral faces; “rh_paracentral_thickness” = para central lobule cortical thickness; “PPI_angry.amy.caudate.thalamus.avg” = average amygdala functional connectivity with the thalamus while watching angry faces vs. shapes; “Total CAPS” = Total scores of CAPS-4.

To further characterize patients within each cluster according to the identified biomarkers, we built classification trees (see Fig. 3). Results showed that left hemisphere EC volume had the greatest influence on clustering - 70 out of 101 participants had left EC volume equal to or greater than 1449mm^3^, out of which there was almost an equal distribution between the two clusters (56%/44%). The other 31 participants had a left EC volume smaller than 1449mm^3^, out of which the vast majority (84%) belonged to HiClus; indicating that subjects with lower EC volume were more likely to belong to the HiClus. Further down on the left branch of the tree, HiClus subjects had larger left caudal middle frontal gyri volume. Down the right branch of the tree, high executive functioning was more associated with LoClus, and vice versa. Further down, low supramarginal gyrus cortical thickness together with high paracentral volume were related to LoClus, while low executive functions with low functional connectivity between the amygdala and the left insula while watching fearful faces vs. shapes was strongly related to HiClus.

**Figure 3.**
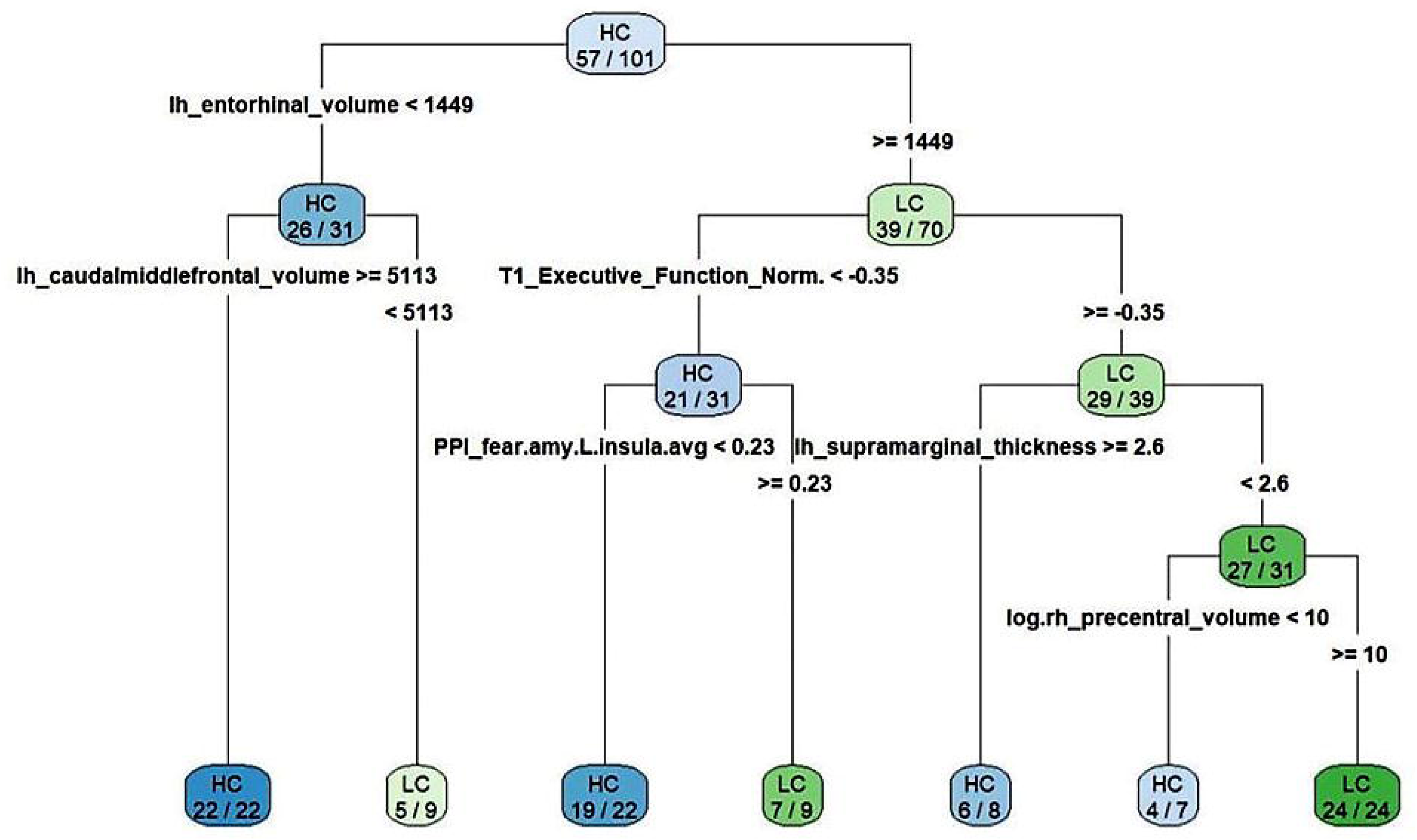
Classification Tree based on the Two Clusters. The classification tree depicts variables important for the division of the participants to the two clusters, starting from the most important one at the top of the tree (left entorhinal cortex volume). Each block in tree is labeled either HC or LC, indicating whether most of the subjects in that block belong to the HiClus or the LoClus (colored bluer or greener, respectively). Furthermore, each block has the number of subjects belonging to the dominant cluster (either HC or LC), and the total number of subjects in that specific block. Inspecting the top block for example, 70 out of 101 participants had left EC volume equal to or greater than 1449mm^3^, out of which 39 belonged to the LoClus. The other 31 participants had a left EC volume smaller than 1449mm^3^, out of which the most (n=26) belonged to the HiClus. For full description please see eResults. “lh_entorhinal_volume” = left entorhinal cortex volume; “T1_Executive_Function_Norm” = executive function score; “lh_caudalmiddlefrontal_volume” = left caudal middle frontal gyrus volume; “lh_supramarginal_ thickness” = left supramarginal gyrus cortical thickness; “PPI_fear.amy.L.insula.avg” = average amygdala functional connectivity with the left insula while watching fearful faces vs. shapes; “log.rh_precentral_volume” = right precentral gyrus volume (after log transformation to create symmetrical distribution).

Finally, a marginal one-way ANOVA was conducted on the potential biomarkers with HiClus/LoClus as the dependent variable. Benjamini-Hochberg adjustment to control the FDR at level 0.05, yielded no significant difference for any potential neurocognitive biomarker between HiClus/LoClus, FDR-corrected (partly due to hundreds of p-values that were adjusted).

To assess robustness and consistency within this sample, the procedure detailed above was repeated with cross-validation. The 3C method was performed on a different percentage of the total subjects (P=20%, 30%, …, 90% of all participants). For each percentage of subjects (P), n=1000 iterations were performed. In each iteration, the 3C methodology was executed on a randomly selected subset of individuals, resulting in different clusters and classification tress. The remaining individuals (which were not included in this iteration), were classified to the new clusters based on the created decision trees. For each P, the mean percentage (and SD) of individuals correctly classified (according to the results of the 3C methodology based on all subjects, see Table 2). For example, when the 3C algorithm was based on only P=20% of the subjects, 83% of the remaining individuals were correctly classified on average.

**Table 2.** Cross-Validation Results. The table present the mean percentage and standard deviation (SD) of subjects which were correctly classified (according to the results of the 3C methodology based on all subjects). For each P, n=1,000 iterations were performed.

| **P** | **20%** | **30%** | **40%** | **50%** | **60%** | **70%** | **80%** | **90%** |
| --- | --- | --- | --- | --- | --- | --- | --- | --- |
| **Mean** | 83% | 86% | 86% | 88% | 89% | 90% | 90% | 95% |
| **SD** | 10% | 7% | 10% | 9% | 7% | 7% | 6% | 7% |

## Discussion

This work illustrates the efficacy of a novel 3-staged (3C) data analytic approach in classifying recent trauma survivors to different “disease load” using a set of potential neurocognitive biomarkers. The 3C ‘hybrid’ data- and theory-driven analytic approach combined current diagnostic-based *categorization*, symptom severity-based *clustering* and the data-driven retrieval of *classifying* using *potential biomarkers*. Unlike supervised machine learning models that ‘flatten’ information and dimensions into one data matrix, the 3C method uniquely and in a stepwise manner combines current PTSD diagnostics with two layers of data-driven exploration: clinical symptom severity and concurrently-recorded objective measurements of cognitive functioning, brain structure and function. It does not impose an extraneous construct (PTSD diagnosis) as an outcome of interest, but rather focus on the actual clustering and richness of phenotypical features associated with post-traumatic psychopathology, thereby may able to create symptom clusters that more reliably represent phenotypical subclasses of theoretical and clinical interest. The biomarkers revealed by the 3C approach are in line with previously documented neural and cognitive correlates of PTSD. Thus, they provide a template for the early objective mechanism-based categorization of trauma survivor psychopathology.

### Potential neurocognitive biomarkers for PTSD

The potential neurocognitive biomarkers for PTSD severity included features obtained from both neuroimaging and cognitive testing. These are not yet proven PTSD biomarkers, as they did not underwent analytical and clinical validations^55^, but rather serve as potential biomarkers from neurocognitive domains. The structural brain features that had the most influence on PTSD classification was entorhinal cortex (EC) volume (both according to importance analysis and classification trees, see Fig. 2 and 3); lower EC volume was associated with higher post-traumatic stress symptom severity. The EC plays an important role in memory, a key feature of post-traumatic psychopathology, as uncontrolled recall of the traumatic event determines symptom severity^56–58^. Another structural feature of importance to classification was the volume of the rostral anterior cingulate cortex (rACC); lower rACC volume was associated with higher PTSD severity. Indeed, rACC volume has previously been associated with PTSD^21,59,60^, and shown to predict cognitive behavioral treatment response for individuals suffering from this disorde^61^; suggesting its potential as a guide for early mechanism-based intervention.

To note, our classifier did not identify several structural abnormalities found in previous cross-sectional PTSD studies^5,13,21,25^. This includes the most replicated finding of small hippocampus volume^62^, but also abnormal amygdala volume, insular cortex, medial and dorsal prefrontal cortices (mPFC and dlPFC respectively)^16,19,20,63–65^. This could stem from our classifier identifying early stage severity-related (or "disease load") biomarkers, rather than DSM-based categories of PTSD diagnosis. Our approach identified structural abnormalities within one-month after trauma as related to symptom severity; since major changes in gray matter volume within this time frame are less likely to occur^66^, these abnormalities might be early predisposing risk factors for chronic PTSD development. Future studies on populations prone to trauma exposure with longitudinal measurement could shed more light on the causal inference of our findings^67–69^.

Functional neuroimaging found amygdala functional connectivity with both the insula and the thalamus to be particularly important for classification (both according to importance analysis and classification trees, see Fig. 2 and 3). Aberrant connectivity of the amygdala with other structures is consistent with previous studies, and abnormal amygdala activation had been hypothesized to contribute to PTSD pathophysiology^70–72^. Moreover, thalamic dysfunction has been found in patients with PTSD, suggesting its role in the disorder’s psychopathology^73,74^. Therefore, although neuroimaging studies have implicated several functional brain abnormalities in the pathophysiology of PTSD, our computational analysis showed that some of these abnormalities are involved in the PTSD severity subtype at the early aftermath of the trauma.

The most significant cognitive factors were cognitive flexibility (according to importance analysis, see Fig. 2) and executive function (according to classification trees, see Fig. 3). Indeed, meta-analyses regarding the role of cognitive functions in PTSD consistently show an impaired ability in executive functioning, including cognitive flexibility among PTSD patients^75,76^. Cognitive flexibility; the ability to switch between two different tasks or strategies^75^ is of particular interest as it has recently been found to correlate with PTSD trajectories in two independent longitudinal cohorts^12^. Specifically, lower scores of cognitive flexibility correlated with increased PTSD symptoms one month after trauma and predicted persistent symptoms one year later. Moreover, disturbances in cognitive flexibility was ameliorated by a cognitive intervention and associated with better treatment outcomes^12^, implying its role as an early recovery related process. It is indeed possible that intact cognitive flexibility enables the individual to better differentiate between threat-related and neutral situations, thus assisting in the extinction of fear-motivated learning, a core-element in PTSD recovery^77^.

### Methodological considerations

Our 3C approach revealed two PTSD subtypes (classes) in recent trauma survivors, correlated with high and low clinical severity, according to total CAPS scores, and across all CAPS subscales (re-experiencing, avoidance, negative alterations in cognitions and mood, and alternations in arousal and reactivity). Our analysis did not find classes representing different clinical subtypes (e.g. dissociative subtype, greater avoidance, etc.). This could be because 3C clustering is based on a given set of clinical measures, and classification is thus based on predetermined measures of potential biomarkers. A larger dataset (i.e., more measures and more participants), could allow for the identification of unique clinical classes of different symptomatic subtypes (possibly more than two). Indeed, in an effort to account for the heterogeneity in PTSD expression, several studies attempted to characterize different subtypes of the disorder. For example, an externalizing subtype characterized by low constraint and high negative emotionality, compared to an internalizing cluster with high negative emotionality and low positive emotionality^78–80^. Another example is a dissociative subtype for patients with PTSD and de-personalization and/or de-realization symptoms, introduced by the fifth edition of the DSM^81–83^. Note that there is no definitive ad-hoc threshold for selection of clinical measurements, different measures strongly efffect the clusters created and the derived classification of potential biomarkers. Lastly, adding longitudinal measures from different time-points following trauma may reveal classes corresponding to PTSD clinical trajectories. This may be cruical for the identification of individuals at risk for developing PTSD, as well as providing appropriate early-stage treatment.

### Conclusion

Our study implemented an innovative computational approach that unveiled novel variables correlated with the morbidity classification of recent trauma survivors, associated with the mechanisms underlying post-traumatic stress symptoms. The method utilized current DSM-based PTSD diagnostic categories and other clinical severity measures of depression and anxiety, as well as a data-driven classification of neurocognitive potential biomarkers. Our results point to an alternative approach for identifying objective variables linked to PTSD severity subtypes (high and low), based on testing within a single session closely after exposure to trauma. If successful, this objective computational classification of potential neurocognitive biomarkers may further guide mechanism-driven interventions for PTSD (e.g. cognitive remediation or neuromodulation treatments). If performed on a larger data set and with more clinical measures and potential biomarkeres, this computational approach may further refine post-traumatic diagnostic subtypes, playing an important role in the treatment management of recent trauma survivors.

## Acknowledgments

This work was supported by award number R01-MH-103287 from the National Institute of Mental Health (NIMH) given to AS (PI), IL and TH (co-Investigators, subcontractors), and had undergone critical review by the NIMH Adult Psychopathology and Disorders of Aging study section. This work was also supported by funding to the Human Brain Project from the European Union Seventh Framework Program (FP7/2007-2013) under grant agreement no. 604102 (HBP). Sagol School of Neuroscience at Tel-Aviv University and the HBP supported ZB fellowships, and Sagol Brain Institute at Tel-Aviv Sourasky Medical Center supported NK and HS fellowships. The authors would like to thank the research team at Tel-Aviv Sourasky Medical Center - including Nili Green, Mor Halevi, Sheli Luvton, Yael Shavit, Olga Nevenchannaya, Iris Rashap, Efrat Routledge and Ophir Leshets - for their major contribution in carrying out this research, including subjects’ recruitment and screening, and performing clinical, cognitive and neural assessments. We also extend our gratitude to all the participants of this study, which completed all the assessments at three different time-points after experiencing a traumatic event.

## Conflict of Interest

The authors declare that they have no financial disclosures and no conflict of interests.

## Trial Registration

Neurobehavioral Moderators of Post-traumatic Disease Trajectories. ClinicalTrials.gov registration number: NCT03756545. https://clinicaltrials.gov/ct2/show/NCT03756545

## References

1. Galatzer-Levy IR, Karstoft KI, Statnikov A, Shalev AY. Quantitative forecasting of PTSD from early trauma responses: A Machine Learning application. J Psychiatr Res. 2014;59:68–76. doi:10.1016/j.jpsychires.2014.08.017

2. Kessler RC, Rose S, Koenen KC, et al. How well can post-traumatic stress disorder be predicted from pre-trauma risk factors? An exploratory study in the WHO World Mental Health Surveys. World Psychiatry. 2014;13(3):265–274. doi:10.1002/wps.20150

3. Perkonigg A, Pfister H, Stein MB, et al. Longitudinal course of posttraumatic stress disorder and posttraumatic stress disorder symptoms in a community sample of adolescents and young adults. Am J Psychiatry. 2005;162(7):1320–1327. doi:10.1176/appi.ajp.162.7.1320

4. Keane TM, Kaloupek DG. Comorbid Psychiatric Disorders in PTSD. Ann N Y Acad Sci. 1997;821(1 Psychobiology):24–34. doi:10.1111/j.1749-6632.1997.tb48266.x

5. Shalev A, Liberzon I, Marmar C. Post-Traumatic Stress Disorder. N Engl J Med. 2017;376(25):2459–2469. doi:10.1056/NEJMra1612499

6. Hyman SE. Can neuroscience be integrated into the DSM-V. Nat Rev Neurosci. 2007;8(9):725–732. doi:10.1038/nrn2218

7. Galatzer-Levy IR, Bryant RA. 636,120 Ways to Have Posttraumatic Stress Disorder. Perspect Psychol Sci. 2013;8(6):651–662. doi:10.1177/1745691613504115

8. Bryant RA, O’Donnell ML, Creamer M, McFarlane AC, Silove D. A multisite analysis of the fluctuating course of posttraumatic stress disorder. JAMA Psychiatry. 2013;70(8):839–846. doi:10.1001/jamapsychiatry.2013.1137

9. Bisson JI, Roberts NP, Kitchiner NJ, Kenardy J. Systematic review and meta-analysis of multiple-session early interventions following traumatic events. Am J Psychiatry. 2009;166(3):293–301. doi:10.1176/appi.ajp.2008.08040590

10. Schuitevoerder S, Rosen JW, Twamley EW, et al. A meta-analysis of cognitive functioning in older adults with PTSD. J Anxiety Disord. 2013;27(6):550–558. doi:10.1016/j.janxdis.2013.01.001

11. Scott JC, Matt GE, Wrocklage KM, et al. A quantitative meta-analysis of neurocognitive functioning in posttraumatic stress disorder. Psychol Bull. 2015;141(1):105–140. doi:10.1037/a0038039

12. Ben-Zion Z, Fine NB, Keynan NJ, et al. Cognitive Flexibility Predicts PTSD Symptoms: Observational and Interventional Studies. Front Psychiatry. 2018;9:477. doi:10.3389/fpsyt.2018.00477

13. Pitman RK, Rasmusson AM, Koenen KC, et al. Biological studies of post-traumatic stress disorder. Nat Rev Neurosci. 2012;13(11):769–787. doi:10.1038/nrn3339

14. Hayes JP, Hayes SM, Mikedis AM, et al. Quantitative meta-analysis of neural activity in posttraumatic stress disorder. Biol Mood Anxiety Disord. 2012;2(1):9. doi:10.1186/2045-5380-2-9

15. Akiki TJ, Averill CL, Abdallah CG. A Network-Based Neurobiological Model of PTSD: Evidence From Structural and Functional Neuroimaging Studies. Curr Psychiatry Rep. 2017;19(11):81. doi:10.1007/s11920-017-0840-4

16. Bremner JD, Elzinga B, Schmahl C, Vermetten E. Structural and functional plasticity of the human brain in posttraumatic stress disorder. Prog Brain Res. 2007;167:171–186. doi:10.1016/S0079-6123(07)67012-5

17. Wang Z, Neylan TC, Mueller SG, et al. Magnetic resonance imaging of hippocampal subfields in posttraumatic stress disorder. Arch Gen Psychiatry. 2010;67(3):296–303.

18. Smith ME. Bilateral hippocampal volume reduction in adults with post-traumatic stress disorder: A meta-analysis of structural MRI studies. Hippocampus. 2005;15(6):798–807. doi:10.1002/hipo.20102

19. Karl A, Schaefer M, Malta LS, et al. A meta-analysis of structural brain abnormalities in PTSD. Neurosci Biobehav Rev. 2006;30(7):1004–1031. doi:10.1016/j.amp.2006.02.006

20. Shin LM, Liberzon I. The Neurocircuitry of Fear, Stress and Anxiety Disorders. Neuropsychopharmacology. 2010;35(1):169–191. doi:10.1038/npp.2009.83

21. Bolsinger J, Seifritz E, Kleim B, Manoliu A. Neuroimaging Correlates of Resilience to Traumatic Events—A Comprehensive Review. Front Psychiatry. 2018;9:693. doi:10.3389/fpsyt.2018.00693

22. Elzinga BM, Bremner JD. Are the neural substrates of memory the final common pathway in posttraumatic stress disorder (PTSD). J Affect Disord. 2002;70(1):1–17. http://www.ncbi.nlm.nih.gov/pubmed/12113915.

23. Rauch SL, Shin LM, Phelps EA. Neurocircuitry Models of Posttraumatic Stress Disorder and Extinction: Human Neuroimaging Research-Past, Present, and Future. Biol Psychiatry. 2006;60(4):376–382. doi:10.1016/j.biopsych.2006.06.004

24. Liberzon I, Abelson JL. Context Processing and the Neurobiology of Post-Traumatic Stress Disorder. Neuron. 2016;92(1):14–30. doi:10.1016/j.neuron.2016.09.039

25. Admon R, Milad MR, Hendler T. A causal model of post-traumatic stress disorder: disentangling predisposed from acquired neural abnormalities. Trends Cogn Sci. 2013;17(7):337–347. doi:10.1016/J.TICS.2013.05.005

26. Koenen KC, Moffit TE, Poulton R, Martin J, Caspi avshalom. Early childhood factors associated with the development of post-traumatic stress disorder: results from a longitudinal birth cohort. Psychol Med. 2007;37(2):181–192. doi:10.1017/s0033291706009019

27. Hendler T, Admon R. Predisposing Risk Factors for PTSD: Brain Biomarkers. In. Comprehensive Guide to Post-Traumatic Stress Disorder. Cham: Springer International Publishing; 2015:1–12. doi:10.1007/978-3-319-08613-2_64-1

28. Schultebraucks K, Galatzer□Levy IR. Machine Learning for Prediction of Posttraumatic Stress and Resilience Following Trauma: An Overview of Basic Concepts and Recent Advances. J Trauma Stress. 2019:1–11. doi:10.1002/jts.22384

29. Feczko E, Miranda-Dominguez O, Marr M, Graham AM, Nigg JT, Fair DA. The Heterogeneity Problem: Approaches to Identify Psychiatric Subtypes. Trends Cogn Sci. 2019:1–18. doi:10.1016/j.tics.2019.03.009

30. Galili T, Mitelpunkt A, Shachar N, Marcus-Kalish M, Benjamini Y. Categorize, Cluster, And classify: A 3-c strategy for scientific discovery in the medical informatics platform of the human brain project. Lect Notes Comput Sci (including Subser Lect Notes Artif Intell Lect Notes Bioinformatics). 2014;8777:73–86.

31. Ben-Zion Z, Fine NB, Keynan NJ, et al. Neurobehavioral Moderators of Post-Traumatic Stress Disorder Trajectories: Prospective fMRI Study of Recent Trauma Survivors. Eur J Psychotraumatol. 2019;10(1). doi:10.1080/20008198.2019.1683941

32. Weathers FW, Keane TM, Davidson JRT. Clinician-administered PTSD scale: A review of the first ten years of research. Depress Anxiety. 2001;13(3):132–156. doi:10.1002/da.1029

33. Blevins CA, Weathers FW, Davis MT, Witte TK, Domino JL. The Posttraumatic Stress Disorder Checklist for DSM-5 (PCL-5): Development. J Trauma Stress. 2015;28:489–498. doi:10.1002/jts.22059

34. Beck, A. T., Steer, R. A., & Brown GK. Beck depression inventory-II. San Antonio. 1996;78 (2)(2):490–498.

35. Beck A, Epstein N, Brown G, Steer RA. An inventory for measuring clinical anxiety: Psychometric properties. J Consult Clin Psychol. 1988;56(6):893.

36. Busner J, Targum SD. The clinical global impressions scale: applying a research tool in clinical practice. Psychiatry (Edgmont). 2007;4(7):28–37.

37. Silverstein S, Berten S, Olson P, et al. Development and validation of a World-Wide-Web-based neurocognitive assessment battery: WebNeuro. Behav Res Methods. 2007;39(4):940–949. doi:10.3758/BF03192989

38. Fischl B. FreeSurfer. Neuroimage. 2012;62(2):774–781.

39. Reuter M, Rosas H, -BF. Highly accurate inverse consistent registration: a robust approach. Neuroimage. 2010. https://www.sciencedirect.com/science/article/pii/S1053811910009717. Accessed November 12, 2019.

40. Ségonne F, Dale AM, Busa E, et al. A hybrid approach to the skull stripping problem in MRI. Neuroimage. 2004;22(3):1060–1075. doi:10.1016/j.neuroimage.2004.03.032

41. Fischl B, Salat DH, Van Der Kouwe AJW, et al. Sequence-independent segmentation of magnetic resonance images. Neuroimage. 2004;23(SUPPL. 1). doi:10.1016/j.neuroimage.2004.07.016

42. Fischl B, Salat DH, Busa E, et al. Whole brain segmentation: Automated labeling of neuroanatomical structures in the human brain. Neuron. 2002;33(3):341–355. doi:10.1016/S0896-6273(02)00569-X

43. Fischl B, Salat DH, Busa E, et al. Whole Brain Segmentation□: Neurotechnique Automated Labeling of Neuroanatomical Structures in the Human Brain. Neuron. 2002;33:341–355.

44. Friston KJ, Ashburner JT, Kiebel SJ, Nichols TE, Penny WD. Statistical Parametric Mapping: The Analysis of Functional Brain Images. Vol 8.; 2007. doi:10.1016/B978-0-12-372560-8.50052-8

45. Hariri AR, Bookheimer SY, Mazziotta JC. Modulating emotional responses: effects of a neocortical network on the limbic system. Neuroreport. 2000;11(1):43–48. doi:10.1097/00001756-200001170-00009

46. Mitelpunkt A, Galili T, Shachar N, Marcus-Kalish M, Benjamini Y. Categorize, Cluster & Classify - The 3C Strategy Applied to Alzheimer’s Disease as a Case Study. Proc Int Conf Heal Informatics. 2015:566–573. doi:10.5220/0005275705660573

47. Jack CR, Bernstein MA, Fox NC, et al. The Alzheimer’s Disease Neuroimaging Initiative (ADNI): MRI methods. J Magn Reson Imaging. 2008;27(4):685–691. doi:10.1002/jmri.21049

48. Benjamini Y, Hochberg Y. Controlling the False Discovery Rate: A Practical and Powerful Approach to Multiple Testing. J R Stat Soc Ser B. 1995;57(1):289–300. doi:10.1111/j.2517-6161.1995.tb02031.x

49. Tibshirani R, Walther G, Hastie T. Estimating the number of clusters in a data set via the gap statistic. J R Stat Soc Ser B Stat Methodol. 2001;63(2):411–423. doi:10.1111/1467-9868.00293

50. Kodinariya TM, Makwana PR. Review on determining number of Cluster in K-Means Clustering. Int J Adv Res Comput Sci Manag Stud. 2013;1(6). www.ijarcsms.com.

51. Archer KJ, Kimes R V. Empirical characterization of random forest variable importance measures. Comput Stat Data Anal. 2008;52(4):2249–2260. doi:10.1016/j.csda.2007.08.015

52. Shachar N, Mitelpunkt A, Kozlovski T, et al. The Importance of Nonlinear Transformations Use in Medical Data Analysis. JMIR Med informatics. 2018;6(2):e27. doi:10.2196/medinform.7992

53. Maechler M, Rousseeuw P, Struyf A, Hubert M, Hornik K. Cluster Analysis Basics and Extensions. R package version 1.14.4. Cran. 2013. http://cran.r-project.org/web/packages/cluster/index.html.

54. Rousseeuw PJ. Silhouettes: A Graphical Aid to the Interpretation and Validation of Cluster Analysis. Vol 20.; 1987.

55. Zhang L, Li H, Benedek D, Li X, Ursano R. A strategy for the development of biomarker tests for PTSD. Med Hypotheses. 2009;73(3):404–409. doi:10.1016/j.mehy.2009.02.038

56. Rubin DC, Berntsen D, Bohni MK. A Memory-Based Model of Posttraumatic Stress Disorder: Evaluating Basic Assumptions Underlying the PTSD Diagnosis. Psychol Rev. 2008;115(4):985–1011. doi:10.1037/a0013397

57. Schuff N, Zhang Y, Zhan W, et al. Patterns of altered cortical perfusion and diminished subcortical integrity in posttraumatic stress disorder: An MRI study. Neuroimage. 2011;54(SUPPL. 1):S62–S68. doi:10.1016/j.neuroimage.2010.05.024

58. Tsao A, Sugar J, Lu L, et al. Integrating time from experience in the lateral entorhinal cortex. Nature. 2018;561(7721):57–62. doi:10.1038/s41586-018-0459-6

59. Kitayama N, Quinn S, Bremner JD. Smaller volume of anterior cingulate cortex in abuse-related posttraumatic stress disorder. J Affect Disord. 2006;90(2-3):171–174. doi:10.1016/j.jad.2005.11.006

60. Woodward SH, Kaloupek DG, Streeter CC, Martinez C, Schaer M, Eliez S. Decreased anterior cingulate volume in combat-related PTSD. Biol Psychiatry. 2006;59(7):582–587. doi:10.1016/j.biopsych.2005.07.033

61. Bryant RA, Felmingham K, Whitford TJ, et al. Rostral anterior cingulate volume predicts treatment response to cognitive-behavioural therapy for posttraumatic stress disorder. J Psychiatry Neurosci. 2008;33(2):142–146. http://www.ncbi.nlm.nih.gov/pubmed/18330460.

62. Woon FL, Sood S, Hedges DW. Hippocampal volume deficits associated with exposure to psychological trauma and posttraumatic stress disorder in adults: A meta-analysis. Prog Neuro-Psychopharmacology Biol Psychiatry. 2010;34(7):1181–1188. doi:10.1016/j.pnpbp.2010.06.016

63. Ferrari MCF, Busatto GF, McGuire PK, Crippa JAS. Structural magnetic ressonance imaging in anxiety disorders: An update of research findings. Rev Bras Psiquiatr. 2008;30(3):251–264. doi:10.1109/ICCIS.2013.425

64. Gilbertson MW, Shenton ME, Ciszewski A, et al. Smaller hippocampal volume predicts pathologic vulnerability to psychological trauma. Nat Neurosci. 2002;5(11):1242–1247. doi:10.1038/nn958

65. Dedovic K, D’Aguiar C, Pruessner JC. What stress does to your brain: A review of neuroimaging studies. Can J Psychiatry. 2009;54(1):6–15. doi:10.1177/070674370905400104

66. Maclaren J, Han Z, Vos SB, Fischbein N, Bammer R. Reliability of brain volume measurements: A test-retest dataset. Sci Data. 2014;1:140037. doi:10.1038/sdata.2014.37

67. Admon R, Leykin D, Lubin G, et al. Stress-induced reduction in hippocampal volume and connectivity with the ventromedial prefrontal cortex are related to maladaptive responses to stressful military service. Hum Brain Mapp. 2013;34(11):2808–2816. doi:10.1002/hbm.22100

68. Bonne O, Brandes D, Gilboa A, et al. Longitudinal MRI Study of Hippocampal Volume in Trauma Survivors With PTSD. Am J Psychiatry. 2001;158(8):1248–1251. doi:10.1176/appi.ajp.158.8.1248

69. De Bellis MD, Hall J, Boring AM, Frustaci K, Moritz G. A pilot longitudinal study of hippocampal volumes in pediatric maltreatment-related posttraumatic stress disorder. Biol Psychiatry. 2001;50(4):305–309. doi:10.1016/S0006-3223(01)01105-2

70. RK S, AP K, RC W, et al. Neural dysregulation in posttraumatic stress disorder: evidence for disrupted equilibrium between salience and default mode brain networks. Psychosom Med. 2012;74(9):904–911 8p. doi:10.1097/PSY.0b013e318273bf33

71. Rabinak CA, Angstadt M, Welsh RC, et al. Altered amygdala resting-state functional connectivity in post-traumatic stress disorder. Front Psychiatry. 2011;2(NOV):62. doi:10.3389/fpsyt.2011.00062

72. Nicholson AA, Sapru I, Densmore M, et al. Unique insula subregion resting-state functional connectivity with amygdala complexes in posttraumatic stress disorder and its dissociative subtype. Psychiatry Res - Neuroimaging. 2016;250:61–72. doi:10.1016/j.pscychresns.2016.02.002

73. Lanius RA, Williamson PC, Densmore M, et al. Neural correlates of traumatic memories in posttraumatic stress disorder: A functional MRI investigation. Am J Psychiatry. 2001;158(11):1920–1922. doi:10.1176/appi.ajp.158.11.1920

74. Yin Y, Jin C, Hu X, et al. Altered resting-state functional connectivity of thalamus in earthquake-induced posttraumatic stress disorder: A functional magnetic resonance imaging study. Brain Res. 2011;1411:98–107. doi:10.1016/j.brainres.2011.07.016

75. Aupperle RL, Melrose AJ, Stein MB, Paulus MP. Executive function and PTSD: Disengaging from trauma. Neuropharmacology. 2012;62(2):686–694. doi:10.1016/j.neuropharm.2011.02.008

76. Polak AR, Witteveen AB, Reitsma JB, Olff M. The role of executive function in posttraumatic stress disorder: A systematic review. J Affect Disord. 2012;141(1):11–21. doi:10.1016/j.jad.2012.01.001

77. Scott WA. Cognitive Complexity and Cognitive Flexibility. Sociometry. 1962;25(4):405. doi:10.2307/2785779

78. Miller MW, Resick PA. Internalizing and Externalizing Subtypes in Female Sexual Assault Survivors: Implications for the Understanding of Complex PTSD. Behav Ther. 2007;38(1):58–71. doi:10.1016/J.BETH.2006.04.003

79. Miller MW, Greif JL, Smith AA. Multidimensional Personality Questionnaire profiles of veterans with traumatic combat exposure: Externalizing and internalizing subtypes. Psychol Assess. 2003;15(2):205–215. doi:10.1037/1040-3590.15.2.205

80. Miller MW, Kaloupek DG, Dillon AL, Keane TM. Externalizing and internalizing subtypes of combat-related PTSD: A replication and extension using the PSY-5 scales. J Abnorm Psychol. 2004;113(4):636–645. doi:10.1037/0021-843X.113.4.636

81. Lanius RA, Brand B, Vermetten E, Frewen PA, Spiegel D. The dissociative subtype of posttraumatic stress disorder: rationale, clinical and neurobiological evidence, and implications. Depress Anxiety. 2012;29(8):701–708. doi:10.1002/da.21889

82. Lanius RA, Vermetten E, Loewenstein RJ, et al. Emotion modulation in PTSD: Clinical and neurobiological evidence for a dissociative subtype. Am J Psychiatry. 2010;167(6):640–647. doi:10.1176/appi.ajp.2009.09081168

83. Swart S, Wildschut M, Draijer N, Langeland W, Smit JH. Dissociative Subtype of Posttraumatic Stress Disorder or PTSD With Comorbid Dissociative Disorders: Comparative Evaluation of Clinical Profiles. Psychol Trauma Theory, Res Pract Policy. 2019. doi:10.1037/tra0000474

