## Supplemental Materials for "Potential Neurocognitive Biomarkers for Post Traumatic Stress Disorder (PTSD) Severity in Recent Trauma Survivors"

#### Online Only Materials

eMethods

eResults

eReferences

eFigure 1. Difference in PTSD Symptom Severity between Clusters

eTable 1. Top Ten Most Important Potential Biomarkers

**EMethods**

**Participants**

Participants were trauma survivors, admitted to the Tel-Aviv Sourasky Medical Center’s Emergency Room (ER) after exposure to one of the following traumatic experiences: car accidents, work-related car accidents, recurring car accidents, bicycle accidents, assaults, terror attacks, hostilities, drowning, mass casualty incidents, robbery or electrocution. Only individuals aged 18-65 who were able to read and comprehend Hebrew (language used in self-report questionnaires, neurocognitive tests and tasks inside the MRI) were considered for a telephone screening interview.

Eligible individuals were not included in the study if they sustained severe head injury or were in a coma upon ER arrival; had a known medical condition that interfered with their ability to give informed consent, cooperate with screening and/or treatment; were previously diagnosed with PTSD prior to the study; had a lifetime history of psychotic illness, had a substance abuse disorder, expressed suicidal ideations or had any other medical or psychological condition that constitute treatment priority upon potential enrollment. Individuals who were not eligible for an MRI scan were similarly excluded. Prisoners and active service members of the Israel Defense Force (IDF) were not included. In addition, within these criteria, survivors who expressed clinically significant PTSD symptoms during the first month after trauma were given preference of inclusion in the study, as they were at higher risk of developing chronic PTSD.

A total of 2,944 individuals admitted to the ER were contacted by phone and agreed to undergo a telephone screening interview. Of those, 114 individuals met a PTSD diagnosis according to the phone interview (excluding the “one-month duration” criteria), completed clinical and neurocognitive assessments, and underwent a structural and functional MRI scan. Thirteen individuals were excluded from further statistical analysis due to missing data (10% or more missing), with a remaining 101 participants included in the final analysis presented here.

**Clinical Assesment**

The Clinician-Administered PTSD Scale (CAPS) is a structured clinical interview evaluating the four PTSD symptom criteria according to the DSM-IV (CAPS-4) on dimensions of frequency, intensity, and severity ^1^. The CAPS contains explicit, behaviorally anchored questions and a rating scale descriptors to enhance reliability^2^. The CAPS-4 continuous total scores were calculated by summing all of the individual sub-items. The Hebrew version used in this study was cross-translated and compared with the original English instrument.

The PTSD Checklist for DSM-IV (PCL) assessed self-reported PTSD symptoms^3^. The PCL was shown to have high internal consistency, test-retest reliability, convergent and discriminant validity^4^, as well as a high correlation with the CAPS^5^.

The Beck Depression Inventory (BDI) served to evaluate self-reported current depressive symptoms^6^. The BDI was found to be a good predictor of chronic PTSD at one-week and one-month after trauma^7^.

The Beck Anxiety Inventory (BAI) was used as a measure of self-reported anxiety symptoms. The BAI displayed high internal consistency (α=0.92) and good test-retest reliability over one week (r=0.75)^8^.

The Participants’ Clinical Global Impression Scale (CGI-P) evaluated the patients’ subjective impression on a scale of 1 (“normal feeling”) to 7 (“the worst feeling there is”)^9^. The CGI offers an easily understood practical measurement tool^10^.

**Cognitive Functioning**

WebNeuro: An Internet-based, comprehensive battery of neurocognitive functioning, previously validated against traditional neurocognitive tests^11^. To standardize testing conditions, all tests were conducted at our laboratory in Hebrew. Performance on the different tasks was calculated using an automated software program that derived standardized Z-scores for each participant on each of the following eleven neurocognitive domains: motor coordination, processing speed, sustained attention, controlled attention, cognitive flexibility, response inhibition, working memory, recall memory, executive function, emotion identification, and emotional bias. For a full description of the tasks that were administered, please see Ben-Zion et al. (2018)^12^.

**MRI Acquisition and Analysis**

To allow high-resolution structural images a T1-weighted 3D Sagittal MPRAGE pulse sequence (TR/TE = 2400/2.29 ms, flip angle = 8º, voxel size 0.7x0.7x0.7 mm, FOV = 224×224 mm) was used. Functional whole-brain scans were performed in an interleaved bottom-to-top order, using a T2*-weighted echo planar imaging free induction decay (EPI-FID) pulse sequence (TR/TE=2000/28 ms, flip angle=90º, voxel size 3.0x3.0x3.0 mm, FOV = 224×224 mm, slice thickness=3 mm, 36 slices per volume).

Briefly, cortical reconstruction and volumetric segmentation using Freesurfer included: motion correction and the averaging of multiple volumetric T1 weighted images, removal of non-brain tissue using a hybrid watershed/surface deformation procedure, automated Talairach transformation, segmentation of the subcortical white matter and deep gray matter volumetric structures intensity normalization, tessellation of the gray matter-white matter boundary, automated topology correction, and surface deformation following intensity gradients to optimally place the gray/white and gray/cerebrospinal fluid borders at the location where the greatest shift in intensity defines the transition to the other tissue class.

Preprocessing and statistical analysis using SPM 12 included: Slice time correction was performed using one slice before the last as the reference slice. Head motions were corrected by six-parameter rigid body spatial transformations, using three translations and three rotation parameters, with the first image serving as a volume reference. A 4^th^degree interpolation was applied to detect and correct head motions. Functional maps were automatically co-registered to corresponding structural maps using an objective function of normalized mutual information (NMI). The complete dataset was transformed into MNI space and spatially smoothed with an isotropic 6mm full-width at half-maximum (FWHM) Gaussian kernel.

For the Emotional Faces Matching Task, subjects were instructed to select the face/shape (located at the bottom right or bottom left of the screen) that matches the target face/shape (located at the top of the screen), as accurately and as quickly as possible. The tasks included four blocks of shapes (that were used as a baseline) and four blocks of emotional faces (angry, fearful, surprised and neutral faces). The order of the blocks of emotional faces was counterbalanced between subjects using four different versions for this task.

**Procedure**

A member of the research team identified potentially trauma-exposed patients using the ER medical records. Within 10–14 days after potential trauma exposure, and after being discharged from the hospital, these individuals were contacted for an initial telephone screening, which was conducted by MA-level clinicians that were trained in the specific assessment tools (for detailed description, see Fine et al. (2018)^13^). After obtaining verbal consent, the PCL5 was administered to assess the risk of PTSD development. Those who met PTSD symptom criteria (except the “one-month duration” criteria) and did not meet any of the exclusion criteria, received verbal information about the study. Participants were subsequently invited to participate in two meetings, both within 30 days from ER admission: (1) An in-person clinical assessment that included administration of the CAPS, self-report questionnaires (BDI, BAI, PCL, CGI) and the WebNeuro neurocognitive battery; (2) A structural and functional MRI scan. Each meeting took approximately 3 hours (total of 6 hours for both), for which participants received financial remuneration at the end of the assessment, in accordance with the ethics committee regulations and approval.

**Algorithms, Codes and Software**

Data imputation was performed using the 5-nn method in order to deal with missing data (less than 1% of the full dataset). All variables with asymmetrical distributions were then transformed into a symmetrical distribution using “Auto-Neta”^14^. Analyzes were performed using R. Importance was calculated using the {randomForest} R package^15^. Importance was measured as the marginal loss of classification accuracy for each variable by randomly permuting it on the test (out of bag) validation set. For clustering we used the R package {cluster}^16^. The classification and regression tree (CART) were constructed using the {rpart} R package.

**eResults**

**3C Classification Results**

Classification trees were built in order to further characterize the subjects of the two clusters. Inspecting the first split of the tree, results show that left hemisphere entorhinal cortex (EC) volume greatly influenced the clustering *(see first split on Fig. 5).* Seventy out of 101 participants had left EC volume equal to or greater than 1449mm^3^, out of which there was almost an equal distribution between the two clusters (56%/44%). The other 31 participants had a left EC volume smaller than 1449mm^3^, out of which the vast majority (84%) belonged to HiClus; indicating that subjects with lower EC volume were more likely to belong to the HiClus. Further down on the left branch of the tree, HiClus subjects had larger left caudal middle frontal gyri volume. Down the right branch of the tree, high executive functioning was more associated with LoClus, and vice versa. Further down, low supramarginal gyrus cortical thickness together with high paracentral volume were related to LoClus, while low executive functions with low functional connectivity between the amygdala and the left insula while watching fearful faces was strongly related to HiClus.

**eReferences**

1.	Weathers FW, Keane TM, Davidson JRT. Clinician-administered PTSD scale: A review of the first ten years of research. *Depress Anxiety*. 2001;13(3):132-156. doi:10.1002/da.1029

2.	Blake DD, Weathers FW, Nagy LM, et al. The development of a clinician-administered PTSD scale. *J Trauma Stress*. 1995;8(1):75-90. doi:10.1002/jts.2490080106

3.	Weathers FW, Huska JA, Keane TM. The PTSD Checklist-Civilian Version (PCL-C). *Boston, MA Natl Cent for, 1994‏*. 1991.

4.	Blevins CA, Weathers FW, Davis MT, Witte TK, Domino JL. The Posttraumatic Stress Disorder Checklist for DSM-5 (PCL-5): Development. *J Trauma Stress*. 2015;28:489-498. doi:10.1002/jts.22059

5.	Blanchard E. Psychometric properties of the PTSD checklist (PCL). *Behav Res Ther*. 1996;34(8):669-673. doi:10.1016/0005-7967(96)00033-2

6.	Beck, A. T., Steer, R. A., & Brown GK. Beck depression inventory-II‏. *San Antonio*. 1996;78 (2)(2):490-498.

7.	Freedman SA, Brandes D, Peri T, Shalev AY. Predictors of chronic post-traumatic stress disorder: A prospective study‏. *Br J Psychiatry*. 1999;174(04):353-359. doi:10.1192/bjp.174.4.353

8.	Beck A, Epstein N, Brown G, Steer RA. An inventory for measuring clinical anxiety: Psychometric properties‏. *J Consult Clin Psychol*. 1988;56(6):893.

9.	Guy W. CGI. Clinical Global Impressions. *ECDEU Assess Man Psychopharmacol*. 1976:217-222.

10.	Busner J, Targum SD. The clinical global impressions scale: applying a research tool in clinical practice. *Psychiatry (Edgmont)*. 2007;4(7):28-37.

11.	Silverstein S, Berten S, Olson P, et al. Development and validation of a World-Wide-Web-based neurocognitive assessment battery: WebNeuro. *Behav Res Methods*. 2007;39(4):940-949. doi:10.3758/BF03192989

12.	Ben-Zion Z, Fine NB, Keynan NJ, et al. Cognitive Flexibility Predicts PTSD Symptoms: Observational and Interventional Studies. *Front Psychiatry*. 2018;9:477. doi:10.3389/fpsyt.2018.00477

13.	Fine NB, Achituv M, Etkin A, Merin O, Shalev AY. Evaluating web-based cognitive-affective remediation in recent trauma survivors: study rationale and protocol. *Eur J Psychotraumatol*. 2018;9(1):1442602. doi:10.1080/20008198.2018.1442602

14.	Shachar N, Mitelpunkt A, Kozlovski T, et al. The Importance of Nonlinear Transformations Use in Medical Data Analysis. *JMIR Med informatics*. 2018;6(2):e27. doi:10.2196/medinform.7992

15.	Liaw A, Wiener M. *Classification and Regression by RandomForest*. Vol 2.; 2002. doi:10.1023/A:1010933404324

16.	Maechler, M., Rousseeuw, P., Struyf, A., Hubert, M., Hornik K. Cluster Analysis Basics and Extensions. R package version 1.14.4. *CRAN*. 2013.

**eFigure 1.** Difference in PTSD Symptom Severity between Clusters.

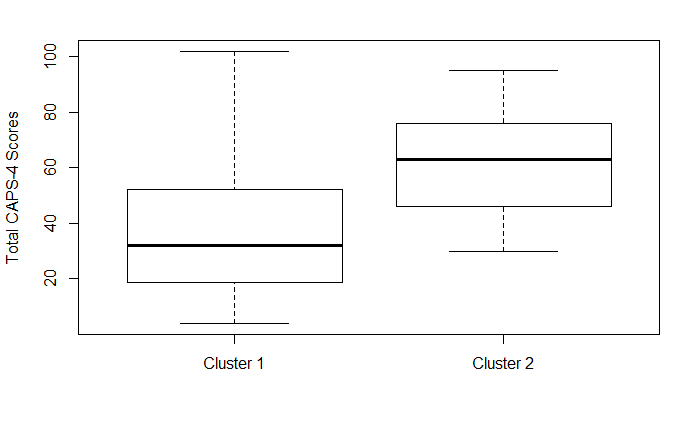
Figure depicts CAPS-4 total scores (Y-axis) for each of the two clusters extracted in stage 2 (clustering).

**eTable 1.** Top Ten Most Important Potential Biomarkers (pBMs)

Table depicts the mean decrease GINI (importance) measure for each pBM. Importance was measured as the marginal loss of classification accuracy for each variable by randomly permuting it on the test (out of bag) validation set.

| **Importance** | **Potential Biomarkers** |
| --- | --- |
| 0.884 | Left entorhinal cortex volume |
| 0.501 | 3^rd^ ventricle volume |
| 0.496 | Average visual area activation while watching neutral faces |
| 0.487 | Cognitive flexibility score |
| 0.473 | Right paracentral lobule cortical thickness |
| 0.441 | 4^th^ ventricle volume |
| 0.429 | Right rostral anterior cingulate cortex (rACC) volume |
| 0.419 | Average connectivity (PPI) between amygdala and thalamus while watching angry faces |
| 0.412 | Left pars-orbitalis volume |
| 0.405 | Right transverse temporal gyrus volume |
